# Deep functional profiling of gene sets using Large Language Models: A blueprint for tailored, context-aware functional annotation

**DOI:** 10.1101/2024.12.12.628275

**Authors:** Taushif Khan, Marina Yurieva, Basirudeen Syed Ahamed Kabeer, Mohammed Toufiq, Darawan Rinchai, Karolina Palucka, Damien Chaussabel

## Abstract

In this study, we present a proof-of-concept approach leveraging Large Language Models (LLMs) for the deep functional profiling of gene sets, addressing the limitations of traditional functional annotation tools. By employing a stepwise prompting strategy with OpenAI’s GPT-4, we systematically retrieve, consolidate, and score immune functions associated with each gene in a module, generating ranked lists of functions with detailed justifications and supporting literature evidence. As a blueprint for this approach, we demonstrate how this in-depth, context-aware analysis can characterize Module M10.4 from the BloodGen3 transcriptional module repertoire, confirming its central role in neutrophil-mediated innate immunity and antimicrobial defense while uncovering a nuanced network of immune functions overlooked by conventional methods. The LLM-based workflow identifies cell types and transcriptional programs driving these functions, offering a more granular and quantitative assessment of the gene set’s functional associations compared to direct LLM prompting or pathway enrichment analysis. Our results showcase the potential of LLMs in providing biologically meaningful interpretations of gene expression signatures, accounting for cellular composition changes and transcriptional regulation. This initial proof-of-concept study establishes a foundation for the automated, high-throughput functional profiling of transcriptional signatures. By integrating advanced language models with traditional literature-based validation, we present a powerful strategy for unraveling the complex biology of gene sets and transcriptional modules, ultimately facilitating the translation of systems-level molecular data into actionable knowledge.

## BACKGROUND

Over the past few decades, large-scale profiling techniques have become pivotal in contemporary biomedical research, marked by the advent of multimodal analyses (multi-omics) and single-cell profiling technologies (1,2). These advancements, alongside a significant reduction in sequencing costs, have made it possible to measure a vast array of parameters simultaneously (3). As a result, these techniques have become increasingly accessible, leading to their widespread use across various fields of biomedical research (4). Such analyses typically yield complex signatures, encompassing tens to thousands of analytes, presenting a significant interpretative challenge (5). This challenge is further compounded by the need to assimilate the extensive, often unstructured literature associated with these gene or analyte collections (6).

To address the complexities of interpreting large-scale profiling data, standardized annotations or definitions for each analyte are indispensable (7). However, traditional tools such as ontology or pathway enrichments, while useful, often provide only a surface-level understanding of the data (8,9). These methods tend to be generic and fail to incorporate the nuanced biological or experimental contexts that are critical for a deeper understanding of gene functions and interactions (10). Consequently, functional interpretation continues to be a rate-limiting step in systems-scale profiling, highlighting the urgent need for innovative approaches capable of comprehensively interpreting and integrating the complex, multi-layered information presented in biological datasets (11).

Advanced Large Language Models (LLMs), such as OpenAI’s GPT-4 and Anthropic’s Claude, have proven instrumental in harnessing vast amounts of information embedded in biomedical texts (12,13). Our previous work which utilized LLMs for knowledge-driven gene prioritization and selection, also illustrates this capability (14). The process involved complex tasks such as identifying functional convergences, scoring candidate genes based on biological and clinical relevance, and fact-checking justifications – all with minimal human intervention. This work highlighted the remarkable efficiency of LLMs, in processing and synthesizing biomedical information, thereby facilitating a more informed and rapid selection of candidate genes. Building upon these findings, our current study explores the potential of LLMs in unraveling functional associations within sets of co-expressed genes, pushing the boundaries of context-sensitive analysis in gene expression data.

Expanding upon these earlier insights, our present investigation explores the capabilities of LLMs in enabling deeper functional profiling through their ability to unravel complex functional associations within sets of co-expressed genes,(15). This exploration aims to advance the scope of context-aware analysis in gene expression data, leveraging the nuanced comprehension of LLMs to reveal deeper biological insights and interactions that conventional methods may overlook (16).

## METHODS

### BloodGen3 Module repertoire

Detailed accounts of the BloodGen3 repertoire’s development have been previously published. In summary, our approach involved analyzing a set of 16 reference datasets that included 985 distinct blood transcriptome profiles, each representing different disease and physiological conditions such as infectious and autoimmune diseases, pregnancy, and organ transplant scenarios. We detected co-clustering patterns which provided the foundation for creating a weighted network. Within this network, we were able to pinpoint highly interconnected networks, or ‘modules’, and further organized these modules into larger groups called ‘aggregates’, informed by the transcript abundance patterns noted across the 16 datasets. This approach facilitated a dual-layered dimension reduction, yielding a more manageable number of variables, either at the level of individual modules (382 variables) or at the aggregated module level (38 variables).

### Large Language Models

“OpenAI’s GPT-4, the latest iteration in the series of Generative Pre-trained Transformers, represents a significant advancement in natural language processing technologies. GPT-4’s core aim is to parse and generate text that is indistinguishable from that authored by humans. It achieves this through sophisticated unsupervised learning algorithms and is trained on an extensive corpus of text from the internet, encompassing a diverse array of linguistic structures and scenarios. This training ensures GPT-4’s proficiency across various domains and its ability to respond to a range of context-specific queries. Notably, its training concluded in April 2023, which marks the cutoff for its knowledge base; hence, it does not assimilate new information post that date, and its responses reflect the dataset available up to that point in time.”

### Direct prompting of the models

<u>PROMPT:</u> Identify functional convergences among this set of genes: [<u>BPI, CEACAM6, CEACAM8, CTSG, DEFA1, DEFA3, DEFA4, ELA2, LOC653600, LOC728358, LOC728358, LOC728358, LTF, MPO, and OLFM4</u>]. Output the results as a table, with the first column indicating the functional convergences and the second column indicating the associated genes.

### LLM stepwise prompting: in depth delineation of functional convergences

In-depth assessment of functional convergences among a set of genes was achieved through a stepwise prompting approach.

#### Step 1: Gene-by-gene retrieval of associated immunological functions, in triplicates

The first step consisted in prompting GPT-4 to retrieve immunological functions for a given gene. To ensure that the list of immunological functions would be comprehensive we ran this prompt in triplicate.

This was carried out in turn for each of the genes belonging to M10.4.

Outputs for all genes from module M10.4 can be explored interactively via a Prezi application (https://prezi.com/view/jeibv59uTXiRngaPxh4W/).

PROMPT 1: *“In an extensive co-expression analysis across 16 human blood transcriptome datasets, we identified 382 transcriptional modules. One gene, [Gene Symbol], showed notable variance in transcript abundance under diverse pathological and physiological conditions. We are concentrating on securing context-aware functional annotations for [Gene Symbol]. For this, identify the immune processes and states [Gene Symbol] is associated with*.

*There is no need to elaborate or provide justifications for those at this point, which can simply be provided as a list*.

*The output should be generated without resorting to an internet search.”*

Notes:

- Here we specified for the model not to base its output on the results of an internet search as we found the findings to generally be superficial and poorly reproducible.

- The prompts are run in triplicates in three separate sessions.

#### Step 2: Consolidate immunological functions retrieved by the three runs

Next a prompt is used to consolidate the functions identified by the three independent runs.

PROMPT 2: *“The following prompt was run multiple times:*

*“In an extensive co-expression analysis across 16 human blood transcriptome datasets, we identified 382 transcriptional modules. One gene, [Gene Symbol] showed notable variance in transcript abundance under diverse pathological and physiological conditions. We are concentrating on securing context-aware functional annotations for [Gene Symbol]. For this, identify the immune processes and states [Gene Symbol] is associated with*.

*There is no need to elaborate or provide justifications for those at this point, which can simply be provided as a list*.

*The output should be generated without resorting to an internet search.”*

*Consolidate the output generated from those multiple runs. Please provide a comprehensive and detailed list ensuring that no association is omitted. After compiling the list, please double-check to confirm that all significant associations are included:*

*[#1 insert the output from PROMPT 1 from the first run, for the gene in question, = OUTPUT1_Rep1[Gene]]*

*[#2 insert the output from PROMPT 1 from the second run, for the gene in question, = OUTPUT1_Rep2[Gene]]*

*[#3 insert the output from PROMPT 1 from the third run, for the gene in question, = OUTPUT1_Rep3[Gene]]”*

Notes:

- Context is provided in PROMPT2 by including the prompt that was used to generate the output that is to be consolidated.

- A follow-on prompt might be used to ask the model to verify its output for completeness. However, we have found that the first output is dependable and using this follow-on prompt does not produce substantial improvements.

#### Step 3: Generate scores for each gene’s associated immunological functions

The next prompt permits to obtain association scores for the immunological functions identified in the previous step for a given gene.

PROMPT 3: *Generate a score indicating the strength of the association between OLFM4 and each of the immunological processes, states or functions listed below. Scoring range is 0-10*.

*Scoring should be generated without calling on analytics or internet searches. A short justification should be provided*.

*The scoring criteria are as follows:*

*0 - No Evidence Found: No scientific studies or data support any association between this gene and the immunological process in question.*

*1-3 - Very Limited Evidence: Sparse or preliminary evidence suggesting an association. This may include small-scale studies, findings with low statistical significance, or early-stage research that indicates a potential but not well-established link*.

*4-6 - Some Evidence: Moderate evidence from several studies or a few larger studies. Findings are somewhat consistent but may have some contradictions or limitations in study design*.

*7-8 - Good Evidence: Strong and consistent evidence from multiple well-conducted studies, including larger-scale and possibly meta-analyses. Evidence indicates a clear association, though minor uncertainties may still exist*.

*9-10 - Strong Evidence: Overwhelming and consistent evidence from a wide range of studies, including large-scale, high-quality research and comprehensive reviews. The association is well- established and widely accepted in the scientific community*.

*[insert the output from PROMPT 2, for the gene in question, = OUTPUT 2_ [Gene]]*

#### Step 4: Organize the compiled information into a structured CSV file

After compiling and scoring the associated immune functions for each gene (Steps 1-3), we instructed GPT-4 to parse this information and organize it in a CSV file format. To avoid overwhelming the model, we ran the prompt separately for each gene, iteratively appending the information to a single CSV file. The prompt used was as follows:

“Create a .csv file to organize the information provided below. The table should have three columns. In the first column, list each immunological function associated. In the second column, include the symbol of the associated gene for each function listed. In the third column, provide the association score. The information to be organized is as follows: [provide output from earlier step, 1 gene at a time]”

We manually verified the accuracy of the final output CSV file, ensuring that all immune functions were extracted and that the cumulative scores matched the expected values. This verification process confirmed GPT-4’s capability in parsing and organizing the information effectively. To further refine the data, we manually collapsed some of the categories before proceeding to the next step. For example, “inflammation,” “regulation of inflammation,” and “inflammatory processes” were combined into a single category: “inflammation.”

#### Step 5: Compute composite scores for each immunological function linked to the gene set

Next, we instruct the model to compute a composite score for the immunological functions identified in the previous step. This is achieved by aggregating the association scores attributed to each gene in the module, a process conducted separately for each immunological function associated with the module. The following prompt is used:

“The .csv file is attached which has three columns. The first column lists immunological functions. The second column includes the symbol of the associated gene for each function listed. The third column includes the association score for each function and gene listed.

Focusing on a specific function, [indicate function of interest], list all associated genes, their association scores and compute an aggregate association score, which is the sum of all association scores for this function.

For those genes that are listed more than once, use only one score (the average of all the scores for that gene).”

#### Step 6: Generate reports for each function

The outputs from steps 6a, 6b, 6c and 6d, which are detailed below, were used to generate a comprehensive report for each immune function individually. This process involved compiling the summary table (step 6a), cell type associations (step 6b), transcriptional program associations (step 6c), functional convergence narrative (step 6d) into a single document for each immune function.

Once a report was completed for a given immune function, the process was repeated for the next immune function associated with at least three genes from the M10.4 gene set (18 in total). This iterative approach allowed for the systematic generation of detailed reports for each relevant immune function, providing a comprehensive understanding of the functional convergences and biological implications of the genes within the M10.4 module.

The content of the reports can be accessed through an interactive plot that was generated as a companion to this manuscript (https://prezi.com/view/jeibv59uTXiRngaPxh4W/).

*Step 6a: generate a summary table with gene symbols, immune functions, association scores, and detailed narratives*.

For the immune function that was singled out in the previous step, and following the computation of composite scores, we instructed the model to generate a summary table that includes the gene symbol, immune function, association score, and a detailed narrative describing its association with the immune function. The prompt used was as follows:

“Generate a table listing in the first column the gene symbol, in the second column the immune function, in the third column the association score and in a third column a detailed narrative describing its association with the immune function. The narrative should be based on current, peer-reviewed scientific knowledge and include specific roles these genes are known to play in the immune system.”

*Step 6b: Identify cell types associated with the gene set in the context of the specific immune function*.

To further contextualize the functional associations of the gene set, for the same immune function, we instructed the model to identify cell types that might be most specifically associated with the genes, considering the patterns of transcript abundance in whole blood across various diseases. The prompt used was as follows:

“This set of genes displayed similarities in patterns of transcript abundance in whole blood across a wide range of diseases. This can be explained by relative changes in the abundance of leukocyte populations. Based on the narratives generated above, describe which cell types this set of genes could be most specifically associated with, in the context of [indicate function of interest].

The output should be organized in a table, with the first column indicating the context (indicate function of interest), the second column indicating the nature of the association (cell type), the third column indicating the associated cell types, along with the symbols of genes which are associated with the cell type, and the fourth column including a justification.

*Step 6c: Identify transcriptional programs associated with the gene set in the context of the specific immune function*.

To further elucidate the underlying mechanisms driving the coordinated expression of the gene set, we instructed the model to identify transcriptional programs that might be most specifically associated with the genes, considering the patterns of transcript abundance in whole blood across various diseases. The prompt used was as follows:

“This set of genes displayed similarities in patterns of transcript abundance in whole blood across a wide range of diseases. This can be explained by coordinated transcriptional regulation.

Based on the narratives generated above, describe which transcriptional programs this set of genes could be most specifically associated with, in the context of [indicate function of interest]. The output should be organized in a table, with the first column indicating the context [indicate function of interest], the second column indicating the nature of the association (transcriptional program), the third column indicating the associated transcriptional programs, along with the symbols of genes which are associated with the transcriptional program, and the fourth column including a justification.

*Step 6d: Generate a narrative describing the functional convergences among the genes in the context of the specific immune function*.

To synthesize the information gathered in the previous steps, we instructed the model to generate a narrative describing the functional convergences observed among the genes in the context of the specific immune function of interest. The prompt used was as follows:

“Next, generate a narrative describing the functional convergences observed among these genes in the context of [indicate immune function of interest]. The style should be direct and technical and should avoid overstatements.”

#### Step 7: Fact-check individual statements and retrieve backing references

The final step involves fact-checking the narrative generated in the previous step and providing backing references for the statements, as would be expected in a research article. To accomplish this, we utilized the Claude 3 language model, using the following prompt:

“Provide backing references for the statements in this paragraph, where expected in a research article. If no suitable references can be found, please flag and edit out the statements in question” Claude 3 was tasked with identifying relevant references to support the statements made in the narrative. If no suitable references could be found for a particular statement, Claude 3 was instructed to flag and edit out the statement in question. The references provided by Claude 3 were then manually checked for factuality and adequacy in justifying or supporting the corresponding statements. This process ensures that the narrative is grounded in current scientific knowledge and that any unsupported claims are removed.

#### Step 8: Generating summary tables using Claude 3

After compiling all 16 reports into a single PDF file, we utilized the Claude 3 language model to generate multiple summary tables, consolidating the findings from the individual reports. The following prompts were used to generate the summary tables:

1. Generating Table 4 - Immune Functions Associated with the M10.4 Gene Set and Their Cumulative Association Scores: “Please generate a table (Table 4) listing all the immune functions investigated in the attached reports, along with their cumulative association scores. The table should have three columns: ‘Immune Function,’ ‘Cumulative Association Score,’ and ‘Associated Genes (Association Score).’ The rows should be ordered in descending order based on the cumulative association scores.”
2. Generating Table 5 - Cumulative and Individual Association Scores of Immune Functions for Each Gene in the M10.4 Gene Set: “Please create a table (Table 5) summarizing the total association scores for each gene in the M10.4 gene set, along with the individual immune functions and their respective association scores. The table should have three columns: ‘Gene,’ ‘Total Association Score,’ and ‘Immune Functions (Score).’ The rows should be ordered in descending order based on the total association scores. Within each cell in the ‘Immune Functions (Score)’ column, the immune functions should also be ordered in descending order based on their individual association scores.”
3. Generating Table 6 - Immune Functions Associated with the M10.4 Gene Set, Their Justifications, and Backing References: “Please generate a table (Table 6) presenting the immune functions associated with the M10.4 gene set, along with summarized justifications and backing references. The table should have three columns: ‘Immune Function,’ ‘Summarized Justification,’ and ‘Backing References.’ The justifications should be based on the narratives included at the end of the reports, and the backing references should be derived from the references provided in the reports.”
4. Generating Table 7 - Cell Type Associations of the M10.4 Gene Set and Their Corresponding Immune Functions: “Please create a table (Table 7) presenting the cell types associated with the genes in the M10.4 set, based on the immune functions identified in the reports. The table should have three columns: ‘Cell Type,’ ‘Associated Immune Functions,’ and ‘Associated Genes.’ The rows should be ordered in descending order according to the number of associated genes.”
5. Generating Table 8 - Transcriptional Programs Associated with the M10.4 Gene Set and Their Corresponding Immune Functions: “Please generate a table (Table 8) presenting the transcriptional programs inferred from the immune functions associated with the M10.4 gene set, as identified in the reports. The table should have three columns: ‘Transcriptional Program,’ ‘Associated Immune Functions,’ and ‘Associated Genes.’ The rows should be ordered in descending order based on the number of associated genes.”

The resulting tables (Tables 4-8) provide a comprehensive overview of the immune functions, cell types, transcriptional programs, and genes associated with the M10.4 gene set, as well as their respective justifications and backing references.

### Interactive Visualization of Functional Profiling Results

An interactive visualization tool was developed using the Prezi presentation platform (https://prezi.com/) to complement the LLM-based functional profiling workflow. An interactive circle plot was created with genes arranged in the outer circle and immune functions represented by color-coded nodes in the inner circles. The color-coding indicates the range of the cumulative association scores for each immune function, while the edges connecting the genes to the immune functions represent the individual association scores. Detailed reports for each immune function, generated through the LLM-based workflow, were embedded within the Prezi presentation. These reports contain comprehensive information on the gene-function associations, including supporting literature evidence. Users can access these reports by clicking on the corresponding immune function nodes within the interactive plot. The Prezi platform was chosen for its ability to create and share interactive presentations online. The platform’s features allowed for the development of a proof-of-concept user interface that enables users to explore the functional profiling results intuitively. Such a user interface would provide a centralized resource for accessing and navigating the comprehensive information generated by the LLM-based workflow.

### Use of LLMs for streamlining manuscript preparation

We explored the use of Claude 3 “Opus” by Anthropic was employed to assist in the preparation of this manuscript. The authors provided Claude 3 with background information, including an initial draft of the manuscript and key findings, to generate content for specific sections such as the results, discussion, and methods. The LLM-generated text was then iteratively refined through a process of author review, additional prompts for targeted improvements, and manual editing to ensure accuracy, clarity, and coherence. Claude 3 also assisted with literature searches and suggesting relevant references based on its knowledge cutoff of August 2023, which were then manually verified by the authors.

## RESULTS

### Functional resolution of fixed transcriptional module repertoires as a use case

In this proof of concept, our objective was to elucidate functional relationships within a module (a co-expressed gene set), from the BloodGen3 transcriptional module repertoire. This repertoire includes 382 modules identified through co-clustering analysis across 16 patient cohorts, incorporating nearly 1,000 transcriptome profiles (refer to Methods & (17)). Notably, this repertoire serves as a standardized framework for data analysis and visualization across multiple projects, including those involving datasets not initially used in its development. The use of such a stable framework allows for accumulated insights into the functional significance of its components over time. The repetitive application of these modules across various studies, spanning years, underscores the value in investing efforts to understand their functional relevance thoroughly. This in turn inspired our current work to establish a methodology for in-depth functional profiling using Large Language Models (LLMs), with the hope of being able to offer deeper and more nuanced insights than traditional approaches. **Figure 1** provides an overview of the BloodGen3 transcriptional module repertoire construction (A.1) and the LLM-enabled functional profiling workflow (A.2) employed in this study.

**Figure 1:**
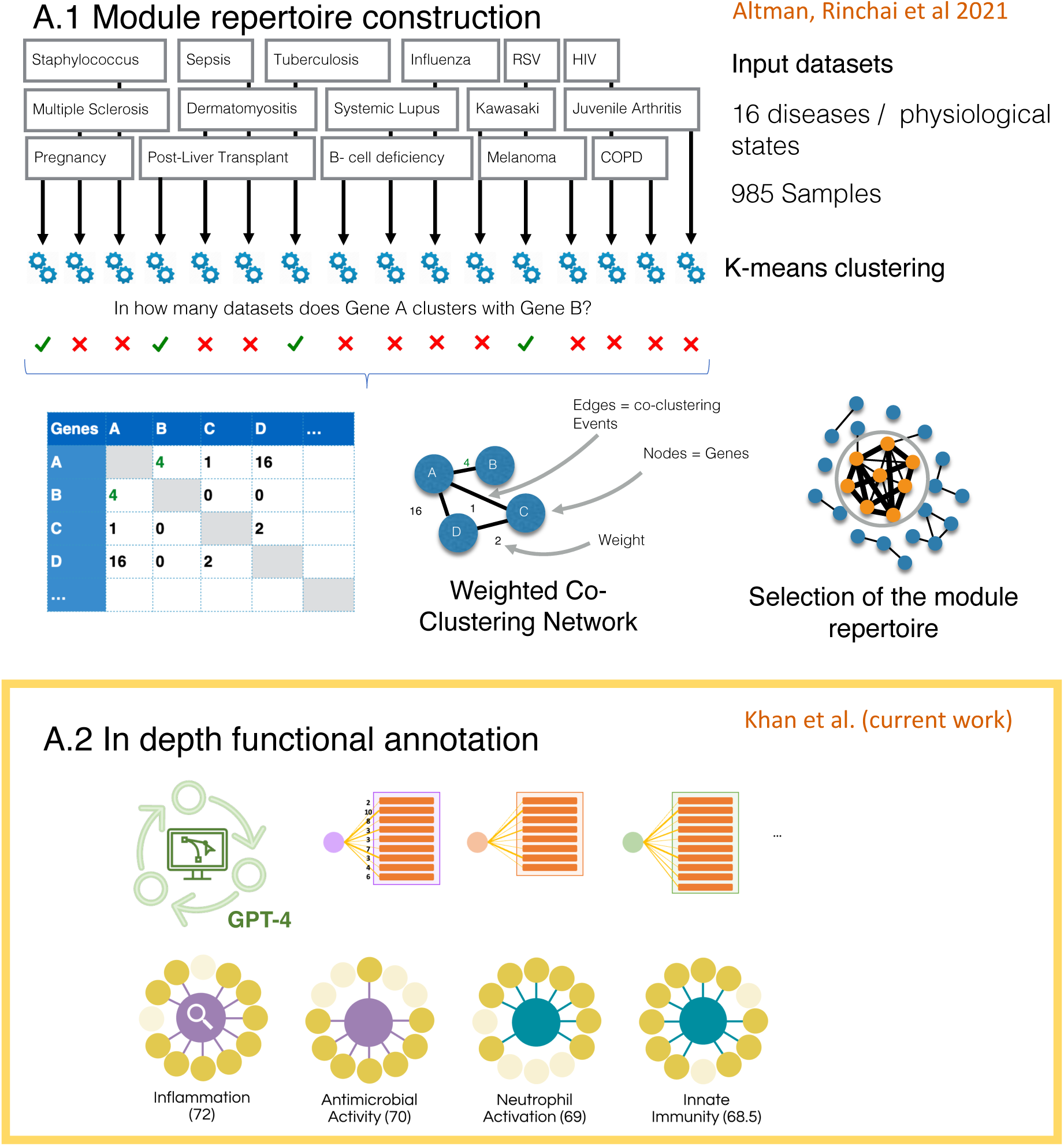
Overview of the development and functional profiling of the fixed BloodGen3 transcriptional module repertoire. (A.1) The construction of the BloodGen3 module repertoire begins with the assembly of an extensive collection of blood transcriptome datasets representing diverse disease states and physiological conditions. These datasets are analyzed to identify co-expression patterns, which form the basis for defining a set of transcriptional modules that constitute a fixed and reusable framework. (A.2) The current work focuses on the in-depth functional profiling of this fixed repertoire using an LLM-based approach. This comprehensive functional characterization is a key step in the development of the BloodGen3 resource, as it enables the interpretation of module-level data across multiple studies and projects over an extended period. The LLM-enabled functional profiling workflow leverages the capabilities of language models to systematically retrieve, consolidate, and score immune functions associated with each module, providing a rich annotation framework to support downstream analysis and interpretation of blood transcriptome data using the BloodGen3 repertoire.

For our initial exploration, we focused on Module M10.4 of the BloodGen3 repertoire, known for its association with neutrophil microbicidal activity, comprising 13 genes: BPI, CEACAM6, CEACAM8, CTSG, DEFA1, DEFA3, DEFA4, ELA2, LOC653600, LOC728358, LTF, MPO, and OLFM4. We evaluated the effectiveness of LLM-enabled workflows against traditional functional profiling tools in identifying functional convergences within this gene set – which is a crucial step for enhancing the utility of fixed, reusable transcriptional module repertoires as platforms for module-level analysis and interpretation.

### Identification of functional associations employing traditional functional profiling tools

Identification of functional associations employing traditional functional profiling tools We utilized Ingenuity Pathway Analysis (IPA), a widely recognized software tool in the scientific community for the functional profiling of gene lists, to analyze the genes comprising Module M10.4. The IPA results revealed several significant canonical pathways associated with Module M10.4, as depicted in Table 1. Notably, ‘Neutrophil degranulation’ and ‘Neutrophil Extracellular Trap Signaling’ pathways were identified with the highest statistical significance, aligning with the known function of the module related to neutrophil activity. The ‘Airway Pathology in Chronic Obstructive Pulmonary Disease,’ ‘Activation of Matrix Metalloproteinases,’ and ‘Antimicrobial peptides’ pathways were also statistically significant, suggesting additional biological contexts where these genes might play crucial roles.

**Table 1:**
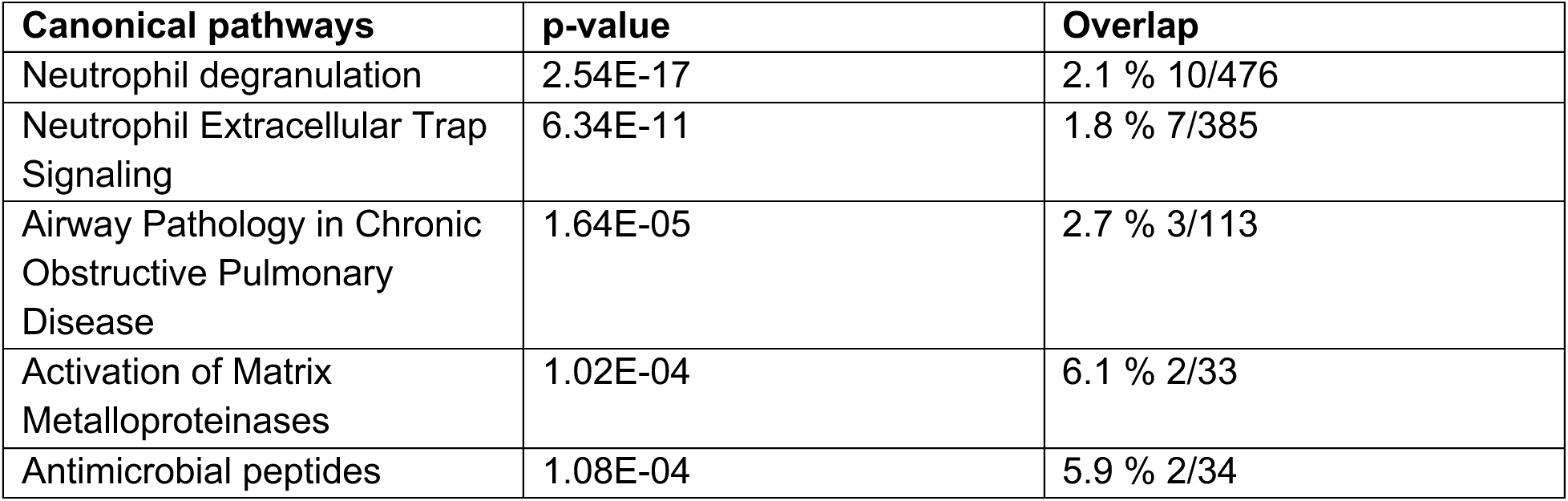
Top IPA canonical pathways for M10.4.

The top IPA diseases and biofunctions, as shown in **Table 2**, support these findings, indicating strong associations with ‘Infectious Diseases,’ ‘Organismal Injury and Abnormalities,’ and ‘Respiratory Disease,’ further substantiating the module’s involvement in antimicrobial and inflammatory responses. The analysis also highlighted several molecular and cellular functions and physiological system development and functions, with ‘Cell Death and Survival,’ ‘Cell-To-Cell Signaling and Interaction,’ and ‘Hematological System Development and Function’ among the most significantly associated functions, reinforcing the relevance of M10.4 genes to key immunological processes.

**Table 2:** Top IPA diseases and biofunction for M10.4.

| Diseases and disorders | p-value range | # Molecules |
| --- | --- | --- |
| Infectious Diseases | 4.03E-02 - 8.78E-21 | 10 |
| Organismal Injury and Abnormalities | 4.50E-02 - 8.78E-21 | 10 |
| Respiratory Disease | 2.59E-02 - 1.09E-12 | 9 |
| Antimicrobial Response | 4.17E-03 - 5.61E-10 | 5 |
| Inflammatory Response | 4.28E-02 - 5.61E-10 | 8 |
| **Molecular and Cellular Functions** |  |  |
| Cell Death and Survival | 2.99E-02 - 2.52E-09 | 7 |
| Cell-To-Cell Signaling and Interaction | 4.24E-02 - 4.36E-07 | 7 |
| Cellular Growth and Proliferation | 3.07E-02 - 1.28E-05 | 4 |
| Cellular Movement | 4.28E-02 - 1.28E-05 | 4 |
| Cell signaling | 2.43E-02 - 6.27E-05 | 4 |
| **Physiological System Development and Function** |  |  |
| Hematological System Development and Function | 4.28E-02 - 4.06E-06 | 6 |
| Immune Cell Trafficking | 4.28E-02 - 6.98E-06 | 6 |
| Organismal Development | 8.33E-03 - 1.28E-05 | 3 |
| Organismal Survival | 1.86E-03 - 6.27E-05 | 3 |
| Tissue Development | 4.64E-02 - 2.85E-04 | 5 |

### Identification of functional associations via direct prompting of the LLMs

Following the functional profiling of the M10.4 gene set with IPA, we proceeded with a direct inquiry approach using GPT-4, as introduced in (14). This method involved a succinct prompt requesting the identification of functional convergences within Module M10.4’s gene set: [BPI, CEACAM6, CEACAM8, CTSG, DEFA1, DEFA3, DEFA4, ELA2, LOC653600, LOC728358, LTF, MPO, OLFM4], with results organized in tabular form. GPT-4 adeptly identified functional themes coherent with the neutrophil activity and antimicrobial function previously known for this module, including Role in Innate Immunity (11 genes), Neutrophil Expression (8 genes), Antimicrobial Activity (6 genes), and Inflammatory Process Involvement (2 genes), with detailed results presented in **Table 3**.

**Table 3:** Functional convergences identified for the genes comprised in module M10.4 through direct prompting of GPT-4.

| Functional Convergence | Associated Genes |
| --- | --- |
| Role in Innate Immunity | BPI, CEACAM6, CEACAM8, CTSG, DEFA1, DEFA3, DEFA4, ELA2, LTF, MPO, OLFM4 |
| Expression in Neutrophils | CEACAM6, CEACAM8, CTSG, DEFA1, DEFA3, DEFA4, ELA2, MPO |
| Antimicrobial Activity | BPI, DEFA1, DEFA3, DEFA4, LTF, MPO |
| Involvement in Inflammatory Processes | CTSG, ELA2 |

The functional associations identified by GPT-4 echoed the significant canonical pathways such as Neutrophil Degranulation and Neutrophil Extracellular Trap Signaling revealed by IPA (detailed in Table 1), underscoring the consistency across both analytical methods. Furthermore, GPT-4’s pinpointing of specific gene functions aligns with the IPA’s broader disease and biofunction categories, like Infectious Disease and Antimicrobial Response (refer to Table 2). This comparison highlights GPT-4’s ability to complement the traditional IPA results by providing a more granular and targeted functional perspective. Notably, while IPA furnished a general overview, GPT-4’s output was remarkable for its specificity, even within broadly defined categories such as Cellular Growth and Proliferation, and Cell Signaling. This specificity is particularly advantageous when delineating the intricate roles of genes within a well-characterized functional context, as evidenced by the findings summarized in **Table 3**.

### Comprehensive Profiling of Immune Functions Associated with the M10.4 Gene Set Using a Stepwise LLM Prompting Strategy

To further enhance the depth and granularity of the functional profiling for the M10.4 gene set, we employed a comprehensive stepwise LLM prompting strategy. A detailed schematic of the LLM-enabled functional profiling workflow, focusing on individual genes and immune functions, is presented in **Figure 2** (detailed in the Methods section). This approach aimed to systematically retrieve, consolidate, and score the immune functions associated with each gene in the module, ultimately generating a ranked list of functions based on their cumulative association scores across the gene set.

**Figure 2:**
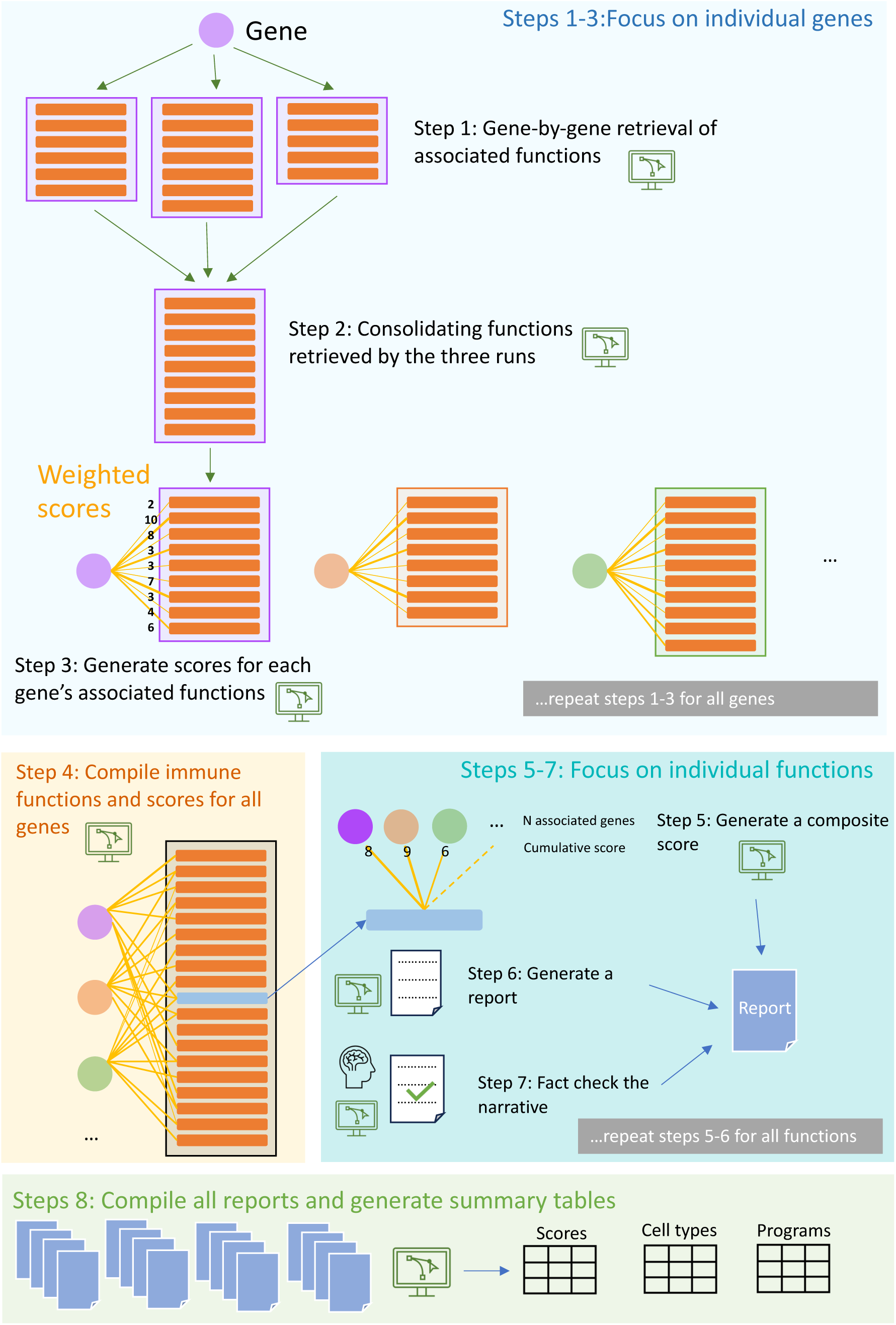
Detailed schematic of the LLM-enabled functional profiling workflow for individual genes and immune functions. The workflow focuses on individual genes in steps 1-3, where associated immune functions are retrieved, consolidated, and scored for each gene in the module. This process is repeated for all genes in the module. In step 4, the immune functions and scores are compiled for all genes. Steps 5-7 focus on individual immune functions, where composite scores are generated, detailed reports are prepared, and the narratives are fact-checked and supported with relevant references. This process is repeated for all immune functions associated with the module. Finally, in step 8, all reports are compiled, and summary tables are generated to provide a comprehensive overview of the module’s functional associations.

The stepwise prompting strategy involved gene-specific retrieval of associated immunological functions, consolidation of functions across multiple runs, scoring of gene-function associations, compilation of module-wide functional landscape, generation of composite scores for each function, and the creation of function-specific narratives. The resulting ranked list of immune functions associated with the M10.4 gene set is presented in **Table 4**. The top-ranking immune functions identified through this approach were Inflammation (cumulative score: 72.0), Antimicrobial Activity (70.0), Neutrophil Activation (69.0), and Innate Immunity (68.5), with sixteen immune functions identified in total with cumulative scores >10. Notably, the stepwise LLM prompting strategy provided a more comprehensive and nuanced understanding of the gene set’s functional associations compared to the traditional IPA analysis (**Tables 1 and 2**) and the direct LLM prompting approach (**Table 3**).

**Table 4:** Immune Functions Associated with the M10.4 Gene Set and Their Cumulative Association Scores. This table presents a comprehensive list of immune functions associated with the genes in the M10.4 gene set, as identified through the LLM-based workflow. The functions are ranked in descending order of their cumulative association scores, which were calculated by summing the individual association scores of each gene linked to a particular function. The table also provides the specific genes associated with each immune function, along with their respective association scores.

| Immune Function | Cumulative Association Score | Associated Genes (Association Score) |
| --- | --- | --- |
| Inflammation | 72 | CEACAM6 (7), CEACAM8 (6), CTSG (7), DEFA1 (7), DEFA1B (7), DEFA3 (7), DEFA4 (6), ELA2 (9), MPO (8), OLFM4 (8) |
| Antimicrobial Activity | 70 | BPI (9), CTSG (8), DEFA1 (10), DEFA1B (9), DEFA3 (9), DEFA4 (9), ELA2 (7), LTF (9) |
| Neutrophil Activation | 69 | BPI (7), CEACAM6 (8), CEACAM8 (9), CTSG (9), DEFA1 (9), ELA2 (10), MPO (9), OLFM4 (8) |
| Innate Immunity | 68 | BPI (7.5), CEACAM6 (7), DEFA1 (9), DEFA1B (10), DEFA4 (7), ELA2 (8), LTF (7), MPO (7), OLFM4 (6) |
| Host Defense | 47.5 | BPI (9), DEFA1 (9), DEFA1B (8), DEFA3 (8), DEFA4 (6), ELA2 (7) |
| Pathogen Clearance | 42 | BPI (7), CEACAM8 (6), CTSG (7), ELA2 (7), MPO (9), OLFM4 (6) |
| Wound Healing | 40 | CEACAM6 (4), CEACAM8 (5), DEFA1 (6), DEFA1B (5), DEFA3 (4), DEFA4 (5), LTF (6), OLFM4 (5) |
| Regulation of Immune Cell Activity | 33 | BPI (5), CEACAM6 (4), CEACAM8 (7), CTSG (6), DEFA1 (5), OLFM4 (6) |
| Response to Infection | 31 | CEACAM8 (8), DEFA1 (8), DEFA1B (7), ELA2 (8) |
| Mucosal Immunity | 29 | CEACAM6 (7), DEFA1 (8), DEFA4 (7), LTF (7) |
| Gastrointestinal Tract Defense | 28 | CEACAM6 (7), DEFA1 (6), DEFA4 (7), OLFM4 (8) |
| Barrier Function | 25 | DEFA1 (7), DEFA1B (8), DEFA3 (4), LTF (6) |
| Chemotaxis | 25 | CTSG (7), DEFA3 (6), DEFA4 (5), ELA2 (7) |
| Autoimmune Diseases | 24 | CEACAM6 (5), CTSG (3), DEFA1B (3), ELA2 (4), LTF (4), OLFM4 (5) |
| Modulation of Immune Response | 22 | CEACAM8 (2), DEFA1 (4), DEFA1B (5), DEFA3 (6), OLFM4 (5) |
| Adaptive Immune Response | 18 | CEACAM6 (5), DEFA1B (4.0), DEFA4 (4.0), LTF (5) |

The application of the stepwise LLM prompting strategy thus not only confirmed the central role of M10.4 in neutrophil-mediated innate immunity and antimicrobial defense but also revealed a more intricate network of immune functions, ranging from Inflammation and Neutrophil Activation to Wound Healing and Adaptive Immune Response. This comprehensive functional profiling enhances our understanding of the biological significance of the M10.4 gene set and demonstrates the utility of leveraging LLMs for in-depth characterization of transcriptional modules.

### Justification and Evidence Supporting the Immune Functions Associated with M10.4

To validate the immune functions identified through the LLM-based approach and ensure that our findings are grounded in existing scientific knowledge, we performed a thorough literature review and compiled a summary of the key evidence supporting each function. This step is crucial for establishing the credibility of our results and providing a solid foundation for the subsequent analyses and interpretations.

Notably, we leveraged Claude 3, to aid in the retrieval of relevant references. By providing Claude 3 with the identified immune functions and their associated genes, we were able to efficiently navigate the vast body of scientific literature and identify the most pertinent studies supporting each function. This innovative application of LLMs in the literature review process not only accelerated the discovery of relevant evidence but also ensured a comprehensive coverage of the available knowledge.

To maintain the highest standards of accuracy and reliability, the relevance of each reference retrieved by Claude 3 was then manually fact-checked. This human-in-the-loop approach permitted to validate the appropriateness of the identified studies and extract the key findings that support the immune functions associated with the M10.4 gene set.

**Table 6** presents the immune functions, their summarized justifications, and the corresponding backing references. The justifications highlight the key mechanisms and pathways through which the genes in the M10.4 set contribute to each immune function, while the backing references provide the necessary support from peer-reviewed literature. The table serves as a comprehensive resource for understanding the biological basis of the identified functions and demonstrates the robustness of the LLM-based approach in capturing meaningful and validated associations.

**Table 5:** Cumulative and Individual Association Scores of Immune Functions for Each Gene in the M10.4 Gene Set. This table presents a summary of the total association scores for each gene in the M10.4 gene set, along with the individual immune functions and their respective association scores. The genes are ranked in descending order based on their total association scores, and within each gene, the immune functions are also listed in descending order of their individual association scores.

| Gene | Total Association Score | Immune Functions (Score) |
| --- | --- | --- |
| DEFA1 | 77 | Antimicrobial Activity (10), Innate Immunity (9), Host Defense (9), Response to Infection (8), Mucosal Immunity (8), Inflammation (7), Barrier Function (7), Gastrointestinal Tract Defense (6), Wound Healing (6), Regulation of Immune Cell Activity (5), Modulation of Immune Response (4) |
| ELA2 | 65 | Neutrophil Activation (10), Inflammation (9), Response to Infection (8), Antimicrobial Activity (7), Host Defense (7), Pathogen Clearance (7), Chemotaxis (7), Regulation of Immune Cell Activity (6), Autoimmune Diseases (4) |
| DEFA1B | 61 | Innate Immunity (10), Antimicrobial Activity (9), Barrier Function (8), Host Defense (8), Inflammation (7), Response to Infection (7), Regulation of Immune Cell Activity (6), Modulation of Immune Response (6), Wound Healing (5), Adaptive Immune Response (4), Autoimmune Diseases (3) |
| OLFM4 | 60 | Inflammation (8), Neutrophil Activation (8), Gastrointestinal Tract Defense (8), Regulation of Immune Cell Activity (6), Innate Immunity (6), Pathogen Clearance (6), Wound Healing (5), Modulation of Immune Response (5), Autoimmune Diseases (5), Adaptive Immune Response (3) |
| DEFA3 | 58 | Antimicrobial Activity (9), Host Defense (8), Inflammation (7), Mucosal Immunity (7), Gastrointestinal Tract Defense (7), Modulation of Immune Response (6), Chemotaxis (6), Barrier Function (4), Wound Healing (4) |
| CEACAM6 | 58 | Neutrophil Activation (8), Inflammation (7), Innate Immunity (7), Mucosal Immunity (7), Gastrointestinal Tract Defense (7), Autoimmune Diseases (5), Adaptive Immune Response (5), Regulation of Immune Cell Activity (4), Wound Healing (4), Barrier Function (4) |
| MPO | 56 | Neutrophil Activation (9), Pathogen Clearance (9), Inflammation (8), Host Defense (8), Response to Infection (8), Antimicrobial Activity (7), Innate Immunity (7) |
| CTSG | 56 | Neutrophil Activation (9), Host Defense (8), Antimicrobial Activity (8), Inflammation (7), Pathogen Clearance (7), Chemotaxis (7), Regulation of Immune Cell Activity (6), Autoimmune Diseases (3) |
| DEFA4 | 56 | Antimicrobial Activity (9), Barrier Function (7), Mucosal Immunity (7), Gastrointestinal Tract Defense (7), Host Defense (7), Innate Immunity (7), Inflammation (6), Regulation of Immune Cell Activity (5), Wound Healing (5), Chemotaxis (5), Modulation of Immune Response (4), Adaptive Immune Response (4) |
| BPI | 56 | Antimicrobial Activity (9), Host Defense (9), Innate Immunity (8), Neutrophil Activation (7), Pathogen Clearance (7), Barrier Function (6), Regulation of Immune Cell Activity (5), Wound Healing (4) |
| LTF | 55 | Antimicrobial Activity (9), Host Defense (7), Mucosal Immunity (7), Gastrointestinal Tract Defense (7), Barrier Function (6), Wound Healing (6), Adaptive Immune Response (5), Autoimmune Diseases (4), Innate Immunity (4) |
| CEACAM8 | 53 | Neutrophil Activation (9), Response to Infection (8), Regulation of Immune Cell Activity (7), Inflammation (6), Pathogen Clearance (6), Chemotaxis (6), Wound Healing (5), Antimicrobial Activity (4), Modulation of Immune Response (2) |

**Table 6.** Immune Functions Associated with the M10.4 Gene Set, Their Justifications, and Backing References. This table presents the immune functions associated with the M10.4 gene set, along with summarized justifications for each function and the corresponding backing references. The justifications and references were compiled through a literature review process aided by the Claude 3 language model and validated by manual fact-checking.

| Immune Function | Summarized Justification | Backing References |
| --- | --- | --- |
| Inflammation | The functional convergence of the genes in the context of inflammation reflects their coordinated roles in antimicrobial defense, immune regulation, and cell signaling. These genes collectively contribute to the immune response by modulating neutrophil functions, producing antimicrobial peptides, and regulating inflammatory signaling pathways. | [1] Medzhitov, R. (2008). Origin and physiological roles of inflammation. *Nature*, 454(7203), 428-435. [2] Nathan, C. (2006). Neutrophils and immunity: challenges and opportunities. *Nature Reviews Immunology*, 6(3), 173-182. [3] Kolaczowska, E., & Kubes, P. (2013). Neutrophil recruitment and function in health and inflammation. *Nature Reviews Immunology*, 13(3), 159-175. |
| Antimicrobial Activity | The genes converge in their roles in the innate immune system's response to microbial threats, showcasing coordinated transcriptional regulation that enhances the antimicrobial capabilities of leukocytes, particularly neutrophils and epithelial cells. The genes encode proteins involved in direct antimicrobial actions, degradation of bacterial proteins, and production of reactive oxygen species. | [1] Hancock, R. E., & Scott, M. G. (2000). The role of antimicrobial peptides in animal defenses. *Proceedings of the National Academy of Sciences*, 97(16), 8856-8861. [2] Ganz, T. (2003). Defensins: antimicrobial peptides of innate immunity. *Nature Reviews Immunology*, 3(9), 710-720. [3] Nguyen, L. T., Haney, E. F., & Vogel, H. J. (2011). The expanding scope of antimicrobial peptide structures and their modes of action. *Trends in Biotechnology*, 29(9), 464-472. |
| Neutrophil Activation | The functional convergence among the genes reflects their integral roles in various aspects of the neutrophil response, including pathogen detection, immune signaling, cell migration, and effector functions. The genes encode proteins involved in microbial killing, cell adhesion processes, proteolytic modification of the extracellular milieu, and modulation of inflammatory responses. | [1] Nauseef, W. M., & Borregaard, N. (2014). Neutrophils at work. *Nature Immunology*, 15(7), 602-611. [2] Amulic, B., et al. (2012). Neutrophil function: from mechanisms to disease. *Annual Review of Immunology*, 30, 459-489. [3] Kolaczowska, E., & Kubes, P. (2013). Neutrophil recruitment and function in health and inflammation. *Nature Reviews Immunology*, 13(3), 159-175. |
| Innate Immunity | The functional convergence of the genes in innate immunity underscores a sophisticated network of defense strategies, encompassing antimicrobial defense, inflammatory regulation, and immune cell recruitment and activation. The genes collectively contribute to the host's ability to rapidly and effectively respond to diverse pathogenic challenges, illustrating the complexity and adaptability of the innate immune system. | [1] Medzhitov, R. (2007). Recognition of microorganisms and activation of the immune response. *Nature*, 449(7164), 819-826. [2] Turvey, S. E., & Broide, D. H. (2010). Innate immunity. *Journal of Allergy and Clinical Immunology*, 125(2), S24-S32. [3] Takeuchi, O., & Akira, S. (2010). Pattern recognition receptors and inflammation. *Cell*, 140(6), 805-820. |
| Host Defense | The functional convergence among the genes in host defense reflects their collective importance in the antimicrobial defense strategy of the host. Through a combination of direct microbial killing and the modulation of immune responses, these genes contribute to the robustness of the host defense mechanisms against a wide array of pathogens, ensuring rapid and effective antimicrobial action. | [1] Hancock, R. E., & Scott, M. G. (2000). The role of antimicrobial peptides in animal defenses. *Proceedings of the National Academy of Sciences*, 97(16), 8856-8861. [2] Ganz, T. (2003). Defensins: antimicrobial peptides of innate immunity. *Nature Reviews Immunology*, 3(9), 710-720. [3] Zasloff, M. (2002). Antimicrobial peptides of multicellular organisms. *Nature*, 415(6870), 389-395. |
| Pathogen Clearance | The functional convergence of the genes in pathogen clearance underscores their collective involvement in critical aspects of the immune response, including antimicrobial defense, immune regulation, and tissue homeostasis. The coordinated expression of these genes likely reflects underlying changes in leukocyte populations and their state of activation, which are central to the pathogenesis and progression of various diseases. | [1] Nauseef, W. M., & Borregaard, N. (2014). Neutrophils at work. *Nature Immunology*, 15(7), 602-611. [2] Rosales, C. (2018). Neutrophil: A cell with many roles in inflammation or several cell types?. *Frontiers in Physiology*, 9, 113. [3] Kolaczowska, E., & Kubes, P. (2013). Neutrophil recruitment and function in health and inflammation. *Nature Reviews Immunology*, 13(3), 159-175. |
| Regulation of Immune Cell Activity | The functional convergence among the genes in the regulation of immune cell activity reflects their roles in antimicrobial defense, modulation of immune responses, and facilitation of cell migration and activation. The genes illustrate a coordinated transcriptional regulation that underpins critical aspects of immune | [1] Turvey, S. E., & Broide, D. H. (2010). Innate immunity. *Journal of Allergy and Clinical Immunology*, 125(2), S24-S32. [2] Medzhitov, R. (2007). Recognition of microorganisms and activation of the immune response. *Nature*, 449(7164), 819-826. [3] Janeway, C. A., & Medzhitov, R. (2002). Innate immune recognition. |
|  | cell activity, ensuring an effective defense against pathogens, the maintenance of tissue integrity, and the regulation of immune cell activity. | Annual Review of Immunology, 20(1), 197-216. |
| Mucosal Immunity | The functional convergence among the genes in mucosal immunity demonstrates a highly coordinated network operating at mucosal surfaces, which not only counters microbial invasion but also maintains the physical and chemical barriers necessary to preserve the health and functionality of mucosal tissues. The genes significantly contribute to the overall resilience and immune competence of the mucosal immune system. | [1] Allaire, J. M., et al. (2018). The Intestinal Epithelium: Central Coordinator of Mucosal Immunity. Trends in Immunology, 39(9), 677-696. [2] Kelsall, B. (2008). Recent progress in understanding the phenotype and function of intestinal dendritic cells and macrophages. Mucosal Immunology, 1(6), 460-469. [3] Okumura, R., & Takeda, K. (2018). Maintenance of intestinal homeostasis by mucosal barriers. Inflammation and Regeneration, 38(1), 5. |
| Wound Healing | The functional convergence of the genes in wound healing underscores a comprehensive transcriptional response geared towards optimizing wound healing outcomes. The coordinated regulation of these genes facilitates a multifaceted approach to wound repair, encompassing pathogen defense, immune regulation, and tissue restoration, which are integral to the successful healing of wounds. | [1] Eming, S. A., et al. (2014). Wound repair and regeneration: mechanisms, signaling, and translation. Science Translational Medicine, 6(265), 265sr6. [2] Gurtner, G. C., et al. (2008). Wound repair and regeneration. Nature, 453(7193), 314-321. [3] Martin, P., & Nunan, R. (2015). Cellular and molecular mechanisms of repair in acute and chronic wound healing. British Journal of Dermatology, 173(2), 370-378. |
| Chemotaxis | The functional convergence among the genes in chemotaxis highlights their coordinated roles in modulating immune responses through specific mechanisms. These genes encode proteins that are fundamentally involved in the recruitment and activation of leukocytes, which are crucial processes during immune responses to infections. | [1] Kolaczowska, E., & Kubes, P. (2013). Neutrophil recruitment and function in health and inflammation. Nature Reviews Immunology, 13(3), 159-175. [2] Nathan, C. (2006). Neutrophils and immunity: challenges and opportunities. Nature Reviews Immunology, 6(3), 173-182. [3] Ley, K., et al. (2007). Getting to the site of inflammation: the leukocyte adhesion cascade updated. Nature Reviews Immunology, 7(9), 678-689. |

By incorporating this additional layer of validation, we aim to strengthen the confidence in our findings and facilitate their integration with existing knowledge in the field. The justifications and evidence presented in **Table 6** not only reinforce the significance of the identified immune functions but also provide valuable context for interpreting the cell type associations and transcriptional programs that underlie these functions. This integrated approach, combining advanced language models with traditional literature-based validation, represents a powerful strategy for unraveling the complex biology of gene sets and transcriptional modules.

### Elucidating Cell Type Associations and Transcriptional Programs Underlying the M10.4 Gene Set’s Immune Functions

To further contextualize the immune functions associated with the M10.4 gene set, we sought to identify the cell types and transcriptional programs that are most likely to be driving these functions. By leveraging the detailed gene-level information generated through the LLM-based approach, we were able to map the genes to their putative cell type associations and infer the transcriptional programs that may be orchestrating their coordinated expression.

**Table 7** presents the cell types associated with the M10.4 gene set, ranked by the number of associated genes. Neutrophils emerged as the most prominent cell type, with 10 out of the 13 genes in the set linked to neutrophil-related functions. This finding reinforces the central role of neutrophils in driving the innate immune response, antimicrobial activity, and inflammatory processes associated with the M10.4 gene set. Various leukocytes and epithelial cells were also identified as important cellular contexts for the expression of these genes, highlighting the involvement of both innate and adaptive immune cells in the gene set’s functional profile.

**Table 7:** Cell Type Associations of the M10.4 Gene Set and Their Corresponding Immune Functions. This table presents the cell types associated with the genes in the M10.4 set, based on the immune functions identified through the LLM-based approach. The cell types are ranked in descending order according to the number of associated genes. The table also lists the specific immune functions linked to each cell type.

| Cell Type | Associated Immune Functions | Associated Genes |
| --- | --- | --- |
| Neutrophils | Inflammation, Antimicrobial Activity, Neutrophil Activation, Innate Immunity, Host Defense, Pathogen Clearance, Regulation of Immune Cell Activity, Response to Infection, Wound Healing, Chemotaxis, Barrier Function, Gastrointestinal Tract Defense, Mucosal Immunity | BPI, CEACAM8, CTSG, DEFA1, DEFA1B, DEFA3, DEFA4, ELA2, MPO, OLFM4 |
| Leukocytes | Antimicrobial Activity, Innate Immunity, Host Defense, Barrier Function, Gastrointestinal Tract Defense | DEFA1, DEFA1B, DEFA3, DEFA4, LTF |
| Epithelial Cells | Inflammation, Innate Immunity, Mucosal Immunity, Gastrointestinal Tract Defense, Wound Healing, Barrier Function | CEACAM6, DEFA1, DEFA4, LTF |
| Monocytes/Macrophages | Regulation of Immune Cell Activity, Wound Healing | OLFM4, LTF |
| Lymphocytes | Adaptive Immune Response | LTF, DEFA4 |
| Immune Response Modulators | Mucosal Immunity, Modulation of Immune Response | DEFA1B |
| Dendritic Cells | Adaptive Immune Response | DEFA1B |
| T-Cells | Adaptive Immune Response | CEACAM6 |
| Natural Killer Cells | Regulation of Immune Cell Activity | CEACAM6 |
| Granulocytes | Response to Infection | CEACAM8 |
| Myeloid Cells | Gastrointestinal Tract Defense | OLFM4 |

**Table 8** summarizes the transcriptional programs inferred from the gene set’s immune functions, again ranked by the number of associated genes. The innate immune response and antimicrobial response programs were found to be the most prominent, aligning with the gene set’s strong associations with innate immunity and host defense. Neutrophil activation and inflammatory response programs were also highly represented, underscoring the importance of these processes in the context of the M10.4 gene set. Notably, the identification of transcriptional programs related to cell adhesion and signaling, immune cell regulation, and lymphocyte modulation suggests that the gene set may also play a role in coordinating the interplay between innate and adaptive immune responses.

**Table 8.** Transcriptional Programs Associated with the M10.4 Gene Set and Their Corresponding Immune Functions. This table presents the transcriptional programs inferred from the immune functions associated with the M10.4 gene set, as identified through the LLM-based approach. The transcriptional programs are ranked in descending order based on the number of associated genes. The table also lists the specific immune functions linked to each transcriptional program.

| **Transcriptional Program** | **Associated Immune Functions** | **Associated Genes** |
| --- | --- | --- |
| Innate Immune Response | Inflammation, Antimicrobial Activity, Innate Immunity, Neutrophil Activation, Host Defense, Pathogen Clearance, Regulation of Immune Cell Activity, Mucosal Immunity, Gastrointestinal Tract Defense, Barrier Function | BPI, DEFA1, DEFA1B, DEFA4, ELA2, LTF, MPO, OLFM4 |
| Antimicrobial Response | Inflammation, Antimicrobial Activity, Innate Immunity, Host Defense, Pathogen Clearance, Mucosal Immunity, Gastrointestinal Tract Defense, Barrier Function, Wound Healing | DEFA1, DEFA1B, DEFA3, DEFA4, ELA2 |
| Neutrophil Activation | Inflammation, Neutrophil Activation, Pathogen Clearance, Response to Infection, Wound Healing, Chemotaxis | CTSG, ELA2, BPI, OLFM4 |
| Inflammatory Response | Inflammation, Antimicrobial Activity, Innate Immunity, Neutrophil Activation, Regulation of Immune Cell Activity, Wound Healing, Gastrointestinal Tract Defense | ELA2, CTSG, LTF, OLFM4 |
| Granulocyte Activation | Response to Infection, Neutrophil Activation, Regulation of Immune Cell Activity | CEACAM8, CTSG, BPI, OLFM4 |
| Pathogen Recognition | Antimicrobial Activity, Innate Immunity, Host Defense, Pathogen Clearance | BPI, LTF, MPO |
| Granule Protein Synthesis | Neutrophil Activation | ELA2, MPO, CTSG |
| Cell Adhesion and Signaling | Inflammation, Neutrophil Activation, Response to Infection, Regulation of Immune Cell Activity, Wound Healing, Adaptive Immune Response | CEACAM6, CEACAM8 |
| Immune Cell Regulation | Regulation of Immune Cell Activity, Wound Healing, Modulation of Immune Response, Adaptive Immune Response | CEACAM6, OLFM4 |
| Immune Cell Activation | Adaptive Immune Response | CEACAM6, DEFA1B |
| Lymphocyte Modulation | Adaptive Immune Response | LTF, DEFA4 |
| Neutrophil Degranulation | Neutrophil Activation | BPI, CTSG, ELA2 |
| Iron Homeostasis | Antimicrobial Activity, Innate Immunity, Host Defense, Mucosal Immunity, Gastrointestinal Tract Defense, Barrier Function, Wound Healing | LTF |
| Endotoxin Neutralization | Antimicrobial Activity, Host Defense, Pathogen Clearance | BPI |

The elucidation of these cell type associations and transcriptional programs provides a more comprehensive understanding of the biological mechanisms underpinning the M10.4 gene set’s immune functions. By identifying the cellular contexts and regulatory pathways that are most relevant to the gene set, we can better appreciate how these genes work in concert to mount effective immune responses. This knowledge can inform future studies aimed at modulating these genes’ expression for therapeutic purposes and guide the development of targeted interventions that harness the innate immune system’s potential in combating disease. Moreover, the approach demonstrated here can be applied to other gene sets or transcriptional modules to unravel the complex interplay between genes, cells, and regulatory programs in the context of immune function and beyond.

### Enhanced Depth and Context-awareness of LLM-based Functional Profiling

A comparative analysis of the results obtained through IPA (Tables 1 and 2) and our LLM-based approach (Table 4) reveals both the complementarity and enhanced depth achieved through the latter. While IPA successfully identified high-level canonical pathways and broad functional categories, our approach provided finer granularity in functional annotation with quantitative association scores. For instance, where IPA identified ‘Neutrophil degranulation’ as a significant pathway (p=2.54E-17), our method dissected this into distinct functional aspects including Neutrophil Activation (score 69), Host Defense (47.5), and Pathogen Clearance (42), each with detailed gene-specific contributions. Beyond this enhanced resolution, our approach provides two critical advantages. First, it generates detailed narratives with backing references (Table 6) for each identified function, offering users a validated knowledge base to deepen their understanding of the underlying biology. Second, and perhaps more significantly, it achieves true context-awareness by explicitly considering factors that specifically affect transcript abundance in whole blood samples. This is evidenced in the systematic mapping of cell type associations (Table 7), which reveals the precise leukocyte populations involved, and the identification of specific transcriptional programs (Table 8) operating within these populations. This context-aware analysis is particularly crucial for blood transcriptomics, where changes in transcript abundance can reflect either altered cellular composition or transcriptional regulation within specific cell populations. Such layered, context-sensitive interpretation represents a significant advance over traditional pathway analysis approaches, providing not just what functions are associated with the gene set, but also how these functions are implemented in the specific context of whole blood.

### Interactive Visualization of the M10.4 Gene Set’s Functional Landscape

The comprehensive functional profiling of the M10.4 gene set using the LLM-based approach revealed a complex network of immune functions associated with the genes in the module. To better visualize and explore these multi-layered associations, we generated an interactive circle plot (**Figure 3 /** https://prezi.com/view/jeibv59uTXiRngaPxh4W/) that integrates multiple elements of our analysis: association scores, detailed narratives with literature backing, cell type associations, and transcriptional programs. This integrated visualization approach permits exploration of the functional landscape at varying levels of granularity.

**Figure 3:**
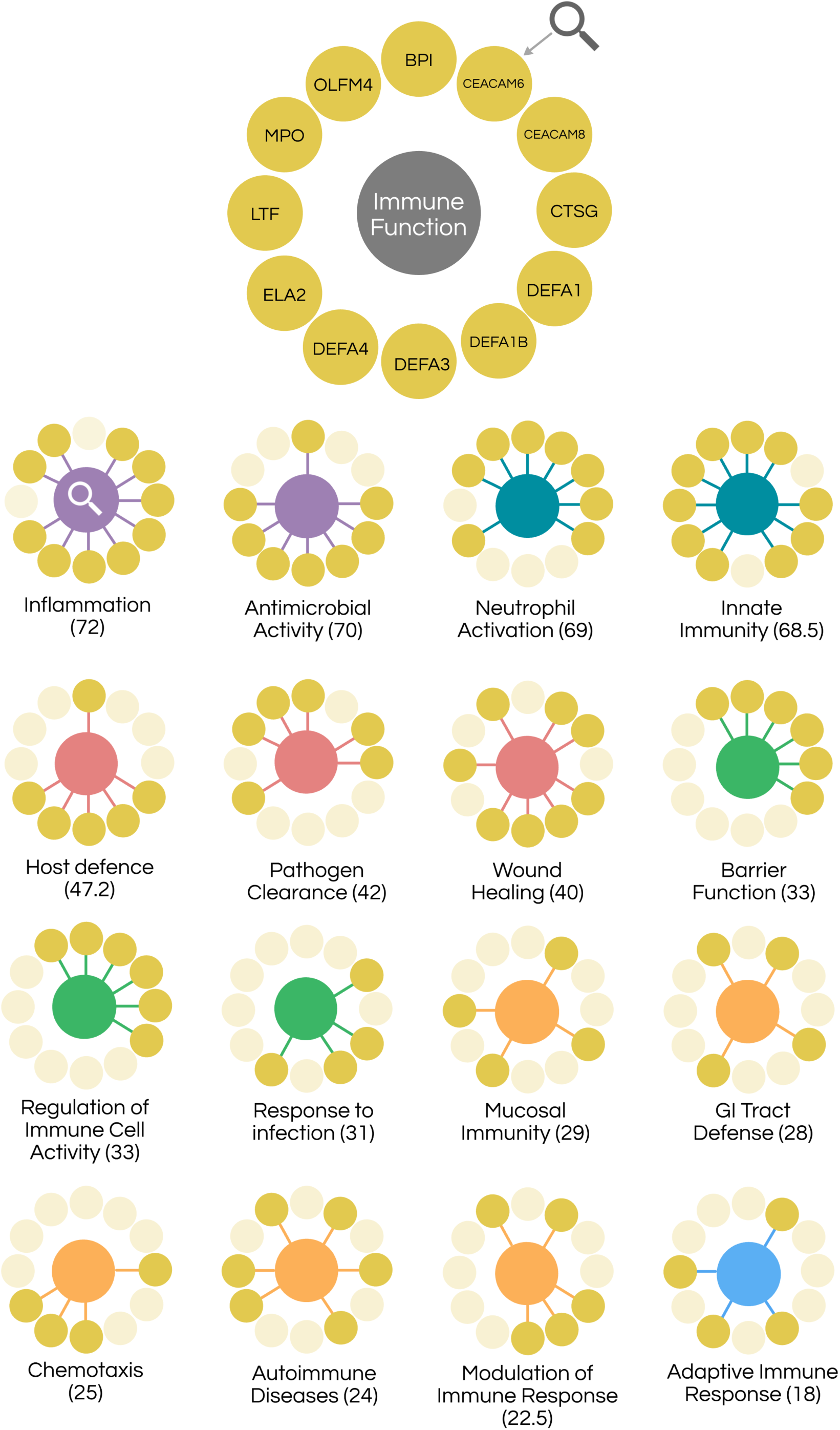
Immune functions associated with the genes in module M10.4 of the BloodGen3 repertoire. The circle plot represents the genes (outer circle) and their associated immune functions (inner circles) identified through the LLM-based functional profiling workflow. The colors of the immune function nodes indicate the range of their cumulative association scores: purple (scores in the 70s), teal (scores in the 60s), red (scores in the 50s), green (scores in the 40s), orange (scores in the 30s), indigo (scores in the 20s), and blue (scores below 20). The edges connecting the genes to the immune functions represent the individual association scores. Detailed reports for each immune function, along with the underlying data, can be accessed interactively via the companion Prezi presentation. (https://prezi.com/view/jeibv59uTXiRngaPxh4W/).

The resulting interactive visualization highlights the intricate relationships between the genes in the M10.4 module and their associated immune functions. The color-coding of the immune function nodes based on their cumulative association scores enables quick identification of the most prominent functions within the module. More importantly, for each functional association, users can access: a) quantitative association scores that indicate strength of evidence, b) detailed narratives explaining the biological mechanisms involved, b) supporting references from peer-reviewed literature, c) relevant cell type contexts, and d) underlying transcriptional programs. This multi-dimensional view provides researchers with both high-level functional insights and deep mechanistic understanding of the M10.4 gene set.

The development of this interactive visualization tool serves as a proof of concept for the potential of combining LLM-based functional profiling with user-friendly visualization techniques. By making multiple layers of functional annotation readily accessible, this approach demonstrates how complex functional profiling data can be made both more interpretable and more deeply informative. With further refinement and adaptation, this methodology could be applied to other gene sets and transcriptional modules, contributing to a better understanding of the functional organization of biological systems.

## DISCUSSION

In this study, we introduced an innovative approach for the functional profiling of gene sets using Large Language Models (LLMs). Our work demonstrates how advanced language models can be leveraged to provide biologically meaningful interpretations of transcriptional data that go beyond traditional annotation approaches. This approach represents a significant step forward in addressing a fundamental challenge in systems biology: the need to extract meaningful biological insights from increasingly complex genomic datasets.

To demonstrate the utility of our approach, we focused on the functional characterization of Module M10.4 from the BloodGen3 transcriptional module repertoire, revealing not only its established role in neutrophil-mediated immunity but also uncovering additional functional aspects through systematic, context-aware analysis. The ability to generate such multi-layered functional interpretations, supported by literature evidence and biological context, showcases the potential of LLMs to enhance our understanding of gene function in specific biological settings. Our findings not only confirmed the central role of Module M10.4 in neutrophil-mediated innate immunity and antimicrobial defense but also revealed a more intricate network of immune functions, including for instance chemotaxis, gastrointestinal tract defense, adaptive immunity and barrier function, which are often overlooked by traditional functional profiling tools. By mapping the genes to their putative cell type associations and inferring the underlying transcriptional programs, our tailored approach also accounted for the fact that changes in transcript abundance in complex cell mixtures, such as blood, are driven by relative changes in cellular composition and the activation or repression of specific transcriptional programs. This context-aware interpretation of gene expression patterns is crucial for uncovering the true biological significance of transcriptional modules and their associated functions.

The application of LLMs for functional profiling of gene sets is a relatively unexplored area, with most existing studies focusing on the use of traditional bioinformatics tools and databases. While these conventional approaches have been valuable in advancing our understanding of gene function and biological pathways, they often rely on predefined annotation categories and may miss important context-specific insights (7,10). In contrast, our LLM-based approach allows for a more flexible and adaptive exploration of gene function, leveraging the vast knowledge encoded in these models to uncover novel and biologically relevant associations. Recent studies have begun to investigate the potential of LLMs and deep learning in various aspects of biological research. For example, Bileschi et al. (2022) demonstrated the utility of deep learning in annotating the protein universe (18), while Grechishnikova (2021) explored the use of transformer neural networks for protein-specific de novo drug generation (19). Our own previous work (Toufiq et al., 2023, (14)) highlighted the ability of LLMs to prioritize candidate genes with minimal human intervention, suggesting the potential of this technology to boost productivity, especially for tasks that require leveraging extensive biomedical knowledge. In the context of viral genomics, Flamholz et al. (2024) showed that protein language models can capture prokaryotic viral protein function, enabling new portions of viral sequence space to be assigned biologically meaningful labels (20). Their work demonstrates the potential of language models to enhance remote homology detection of viral proteins, serving as a useful complement to existing approaches. However, to the best of our knowledge, our current study is the first to apply LLMs specifically for the functional profiling of transcriptional modules in the context of immune function. By demonstrating the ability of LLMs to provide comprehensive and biologically meaningful characterizations of gene sets, our work opens up new avenues for the application of these powerful tools in systems biology and immunology research. Moreover, our approach complements existing methods for the analysis and interpretation of transcriptional module repertoires, such as the BloodGen3 resource used in this study. These repertoires provide a valuable framework for understanding the co-expression patterns and functional relationships among genes across diverse biological conditions (17,21). By integrating LLM-based functional profiling with these established resources, we can gain a more comprehensive and nuanced understanding of the biological processes and pathways associated with specific gene sets, ultimately facilitating the discovery of novel therapeutic targets and biomarkers.

Our study demonstrates the potential of LLMs for comprehensive functional profiling of gene sets, but it is important to acknowledge the limitations of our current approach. The main challenge is the lack of automation in the workflow, which requires significant manual intervention. The stepwise prompting strategy, while effective in eliciting detailed and biologically relevant information from the LLMs, is somewhat intricate and time-consuming. This limits the scalability of the approach, particularly when dealing with larger gene sets or more extensive transcriptional module repertoires.

It is worth noting that the manual corrections required during the fact-checking process were primarily focused on addressing inaccuracies in the references, such as incorrect volumes, page numbers, or journals, or instances where the provided reference did not adequately support a statement. Importantly, we did not encounter instances of reference hallucination when using the specific prompts employed in this study. While hallucinations may still occur with other prompts, even when using advanced models like Claude 3 or GPT-4, the issue has become less prevalent compared to a few months ago.

Furthermore, the investment in time required for the manual curation and fact-checking steps may be justified in the context of our work, which focuses on the functional characterization and interpretation of modules that form a fixed, reusable repertoire. As we have previously demonstrated repertoires can serve as a framework for data analysis and interpretation across numerous studies spanning several years. Thus, the upfront effort invested in thoroughly annotating and validating the functional associations of these modules can yield long-term benefits in terms of the robustness and reproducibility of the analyses.

The proof of concept presented in this study focused on the in-depth characterization of a relatively small gene set, specifically the 13 genes comprising Module M10.4, and their associations with immune functions. However, the workflow can be adapted to retrieve associations between genes and other biological concepts, such as molecular pathways, diseases, or drugs, demonstrating its versatility and potential for a wide range of applications. The applicability of this approach to larger gene sets, containing hundreds or even thousands of genes, remains to be determined, and further investigation is warranted to assess its scalability.

The manual curation and fact-checking steps, which are currently crucial for ensuring the accuracy and reliability of the generated information, may become less resource-intensive as the performance of language models continues to improve. In the preliminary work presented in this manuscript, while some errors were identified in the references provided by the models, these did not lead to any major changes in the gene-immune function associations. This observation suggests that the need for extensive manual fact-checking might decrease over time as the models become more reliable and a sufficient level of trust is established. Nevertheless, it is important to maintain a critical eye and regularly validate the outputs of these models until their performance consistently meets the high standards required for scientific research. It is important to note that our work presents a proof-of-concept workflow, demonstrating the feasibility and potential value of using LLMs for functional profiling of gene sets. However, for this approach to become widely applicable and scalable, further development of tools and automation via API integration is essential.

By addressing these limitations and challenges, future studies can build upon the foundation laid by our work, ultimately enabling the high-throughput, automated functional profiling of large-scale transcriptional module repertoires and other gene set collections. This will facilitate the discovery of novel biological insights and the identification of potential therapeutic targets and biomarkers, accelerating the translation of systems-level molecular profiling data into actionable knowledge.

## DECLARATIONS

### Author Contributions

Conceptualization: DC; Methodology: DC, TK, MY; Investigation: DC, TK, MY; Writing—Original Draft: DC; Writing—Review & Editing: TK, MY, BK, MT, DR, KP, DC; Visualization: DC.

### Conflict of interest/Competing Interests

There are no conflicts to declare.

### Data availability

The BloodGen3 transcriptional module repertoire referenced in this study has been extensively detailed in prior publications (Altman et al., 2021) [17], and the underlying data are available in the NCBI repository under accession ID <u>GSE100150</u>. LLM outputs can be browsed interactively via a Prezi application (https://prezi.com/view/jeibv59uTXiRngaPxh4W/). Reports generated for each associated immune function have been compiled and are available as a supplement (**Supplementary File 1**)

## Notes

### Competing Interest Statement

The authors have declared no competing interest.

## REFERENCES

1. Hasin Y, Seldin M, Lusis A. Multi-omics approaches to disease. Genome Biol. 2017 May 5;18(1):83.

2. Hwang B, Lee JH, Bang D. Single-cell RNA sequencing technologies and bioinformatics pipelines. Exp Mol Med. 2018 Aug 7;50(8):1–14.

3. DNA Sequencing Costs: Data [Internet]. [cited 2024 May 29]. Available from: https://www.genome.gov/about-genomics/fact-sheets/DNA-Sequencing-Costs-Data

4. Manzoni C, Kia DA, Vandrovcova J, Hardy J, Wood NW, Lewis PA, et al. Genome, transcriptome and proteome: the rise of omics data and their integration in biomedical sciences. Brief Bioinform. 2018 Mar 1;19(2):286–302.

5. Subramanian I, Verma S, Kumar S, Jere A, Anamika K. Multi-omics Data Integration, Interpretation, and Its Application. Bioinforma Biol Insights. 2020;14:1177932219899051.

6. Jensen LJ, Saric J, Bork P. Literature mining for the biologist: from information retrieval to biological discovery. Nat Rev Genet. 2006 Feb;7(2):119–29.

7. Khatri P, Sirota M, Butte AJ. Ten years of pathway analysis: current approaches and outstanding challenges. PLoS Comput Biol. 2012;8(2):e1002375.

8. Wadi L, Meyer M, Weiser J, Stein LD, Reimand J. Impact of outdated gene annotations on pathway enrichment analysis. Nat Methods. 2016 Aug 30;13(9):705–6.

9. Haynes WA, Tomczak A, Khatri P. Gene annotation bias impedes biomedical research. Sci Rep. 2018 Jan 22;8(1):1362.

10. Tipney H, Hunter L. An introduction to effective use of enrichment analysis software. Hum Genomics. 2010 Feb;4(3):202–6.

11. Bzdok D, Altman N, Krzywinski M. Statistics versus machine learning. Nat Methods. 2018 Apr;15(4):233–4.

12. Brown T, Mann B, Ryder N, Subbiah M, Kaplan JD, Dhariwal P, et al. Language Models are Few-Shot Learners. In: Advances in Neural Information Processing Systems [Internet]. Curran Associates, Inc.; 2020 [cited 2024 May 29]. p. 1877–901. Available from: https://papers.nips.cc/paper/2020/hash/1457c0d6bfcb4967418bfb8ac142f64a-Abstract.html

13. Deng L, Wang T, Yangzhang null, Zhai Z, Tao W, Li J, et al. Evaluation of large language models in breast cancer clinical scenarios: a comparative analysis based on ChatGPT-3.5, ChatGPT-4.0, and Claude2. Int J Surg Lond Engl. 2024 Apr 1;110(4):1941–50.

14. Toufiq M, Rinchai D, Bettacchioli E, Kabeer BSA, Khan T, Subba B, et al. Harnessing large language models (LLMs) for candidate gene prioritization and selection. J Transl Med. 2023 Oct 16;21(1):728.

15. Eisen MB, Spellman PT, Brown PO, Botstein D. Cluster analysis and display of genome-wide expression patterns. Proc Natl Acad Sci U S A. 1998 Dec 8;95(25):14863–8.

16. Ching T, Himmelstein DS, Beaulieu-Jones BK, Kalinin AA, Do BT, Way GP, et al. Opportunities and obstacles for deep learning in biology and medicine. J R Soc Interface. 2018 Apr;15(141):20170387.

17. Altman MC, Rinchai D, Baldwin N, Toufiq M, Whalen E, Garand M, et al. Development of a fixed module repertoire for the analysis and interpretation of blood transcriptome data. Nat Commun. 2021 Jul 19;12(1):4385.

18. Bileschi ML, Belanger D, Bryant DH, Sanderson T, Carter B, Sculley D, et al. Using deep learning to annotate the protein universe. Nat Biotechnol. 2022 Jun;40(6):932–7.

19. Grechishnikova D. Transformer neural network for protein-specific de novo drug generation as a machine translation problem. Sci Rep. 2021 Jan 11;11(1):321.

20. Flamholz ZN, Biller SJ, Kelly L. Large language models improve annotation of prokaryotic viral proteins. Nat Microbiol. 2024 Feb;9(2):537–49.

21. Chaussabel D, Baldwin N. Democratizing systems immunology with modular transcriptional repertoire analyses. Nat Rev Immunol. 2014 Apr;14(4):271–80.

22. Chaussabel D, Quinn C, Shen J, Patel P, Glaser C, Baldwin N, et al. A modular analysis framework for blood genomics studies: application to systemic lupus erythematosus. Immunity. 2008 Jul 18;29(1):150–64.

